# Unlocking River Biofilm Microbial Diversity: A Comparative Analysis of Sequencing Technologies

**DOI:** 10.1101/2025.06.09.658576

**Authors:** Meri A. J. Anderson, Amy C. Thorpe, Susheel Bhanu Busi, Hyun Soon Gweon, Jonathan Warren, Kerry Walsh, Daniel S. Read

**Author notes:** Corresponding authors: Meri Anderson and Daniel Read.

## Abstract

Freshwater ecosystems are under increasing pressure from pollution, habitat degradation, and climate change, highlighting the need for reliable biomonitoring approaches to assess ecosystem health and identify the causes of biodiversity and ecosystem service loss. Characterisation of freshwater microbiomes has the potential to be an important tool for understanding freshwater ecology, ecosystem health and ecosystem function. High-throughput sequencing technologies, such as Illumina short-read and Pacific Biosciences long-read sequencing, are widely used for microbial community analysis. However, the relative performance of these approaches for monitoring freshwater microbiomes has not been well explored. In this study, we compared the performance of long- and short-read sequencing approaches to assess archaeal and bacterial diversity in 42 river biofilm samples across seven distinct river sites in England by targeting the 16S ribosomal RNA gene.

Our findings demonstrated that longer reads generated by PacBio sequencing provide a higher taxonomic resolution, enabling the classification of taxa that remained unassigned in the short-read Illumina datasets. This enhanced resolution is particularly beneficial for biodiversity assessments because it improves species-level identification, which is crucial for ecological monitoring. Despite this, both sequencing methods produced comparable bacterial community structures regarding taxon relative abundance, suggesting that the sequencing approach does not profoundly affect the comparative assessment of community composition. However, while Illumina offers higher throughput and cost efficiency, PacBio’s ability to resolve complex microbial communities highlights its potential for studies requiring precise taxonomic identification.

## Introduction

DNA sequencing has transformed how we study the living world, opening new opportunities for understanding biodiversity and ecosystem function (Shendure *et al*., 2017; Goodwin *et al*., 2016). This technology has become an increasingly important tool in environmental research, particularly for studying complex multi-kingdom microbial communities (Thompson *et al*., 2017). By revealing the breadth of microbial diversity in environmental samples, DNA sequencing can help researchers and environmental regulators make more informed decisions regarding ecosystem management and conservation (Taberlet *et al*., 2012; Porter & Hajibabaei, 2018). Environmental DNA (eDNA) monitoring has emerged as a transformative method and an essential tool for biomonitoring, with diverse applications, including pathogen detection (Farrell *et al*., 2021), tracking invasive species (Thomas *et al*., 2020), monitoring endangered or cryptic species (Ota *et al*., 2020), assessing biodiversity (Keck *et al*., 2022), and identifying habitat connectivity (Littlefair *et al*., 2023). The effectiveness of these approaches relies on several factors, including the ability to classify and identify DNA markers, including specific taxa of interest, such as pathogens and rare or invasive species, at the highest taxonomic resolution. Accurate and reliable sequencing technologies are pivotal for environmental monitoring because they provide a molecular lens through which researchers can detect and quantify biodiversity, track ecosystem changes and health, monitor invasive species and their potential impacts, evaluate conservation efforts, and unravel complex ecological interactions with unprecedented precision and sensitivity.

Short-read sequencing platforms, such as Illumina, have become widespread owing to their availability, cost-effectiveness and high-throughput capabilities (Bentley *et al*., 2018; Satam *et al*., 2023). The *<*600 bp reads (up to 2x 300 bp) generated by Illumina technology are particularly effective for analysing hypervariable regions of the 16S rRNA gene (Yang *et al*., 2016). However, analyses using these shorter reads can struggle to resolve complex genomic regions and repetitive sequences (van Dijk *et al*., 2018). In contrast, long-read sequencing platforms, such as Pacific Biosciences (PacBio), can generate reads averaging 10-25Kb (Hon *et al*., 2020), offering improved resolution of structural variants and complex genomic regions (Rhoads & Au, 2015; Logsdon *et al*., 2020). Although these longer reads can span multiple repeat and hypervariable regions simultaneously and potentially provide more accurate taxonomic classifications (Callahan *et al*., 2019), they are typically more expensive and have lower throughput (Amarasinghe *et al*., 2020).

The divergent characteristics of long- and short-read sequencing technologies can lead to substantial variations in ecological data derived from molecular assessments. Short-read platforms, such as Illumina, generate high-throughput, precise sequences that excel at detecting abundant taxa but may inadvertently under-represent rare or low-abundance species because of their limited read length and potential amplification biases (Wang *et al*., 2022). Conversely, long-read technologies, such as PacBio and Oxford Nanopore, offer extended genomic fragments that can capture more genetic information, potentially revealing cryptic diversity and providing greater taxonomic resolution (van Dijk *et al*., 2023). These technological differences manifest in ecological assessments through variations in taxonomic detection sensitivity. Short-read methods potentially miss rare ecologically important taxa or underestimate community complexity, while long-read approaches can offer more comprehensive insights into genetic diversity, particularly in complex microbial communities or ecosystems with high genetic heterogeneity. Consequently, choosing a sequencing platform has become a critical methodological consideration that can fundamentally alter the ecological interpretation of molecular biodiversity data.

Previous comparative studies using Illumina and PacBio sequencing technologies have revealed significant variations in methodology and performance across different research contexts. However, most comparative studies have been conducted on model organisms or within well-characterised ecosystems, limiting their applicability to diverse ecological contexts (Ferrarini *et al*., 2013; Zhang *et al*., 2020; Galata *et al*., 2021; Barthélémy *et al*., 2024). Although Gao *et al*. described the utility of long-read data for characterising deep-sea surface sediments (Gao *et al*., 2024), existing comparative analyses often fail to comprehensively address how different sequencing technologies might differentially represent complex ecological interactions and biodiversity gradients. These limitations create a significant research opportunity to develop a more nuanced understanding of how sequencing technology performance varies across different ecological contexts, particularly in under-studied aquatic ecosystems.

Our study compared short-read (Illumina, ca. 235 bp) and long-read (PacBio, ca. 1,600 bp) sequencing to analyse epilithic river biofilm bacterial communities using 16S rRNA gene sequencing. We hypothesised that long-read sequencing would provide greater taxonomic resolution and a more comprehensive understanding of biodiversity than short-read sequencing, particularly for distinguishing closely related bacterial species (Johnson *et al*., 2019; Tedersoo *et al*., 2018). To test this, we sequenced DNA from 42 biofilm samples using short- and long-read sequencing methods to compare community characteristics, including the overlap and uniqueness of bacterial taxa detected using each approach. By addressing the current knowledge gaps in the performance and application of different sequencing approaches, this study enhances our understanding of the optimal use of sequencing technologies for environmental monitoring (Ruppert *et al*., 2019).

## Methods

### Sample Collection

Epilithic biofilm samples were collected from rivers across England as part of the Environment Agency’s River Surveillance Network (RSN) monitoring program, following the standard sampling method described by Kelly *et al*. (2020). The total sampling campaign encompassed 2,101 river biofilm samples collected from 861 sites between 2021 and 2023; a detailed description of the methods for sample collection and short-read sequencing is available in the Environment Agency report (Environment Agency, 2024). This study focused on a subset of 42 samples collected from seven sites twice a year in 2021, 2022, and 2023 (Fig. 2). Land cover maps from UKCEH (Morton *et al.,* 2021) were used to describe dominant land cover types for the upstream catchment of each site (Supp table 3). Samples were obtained by scraping five stones or macrophytes into a tray containing 50 ml of river water. The upper surfaces were brushed with a clean toothbrush to remove biofilms. 5 ml of the biofilm suspension was removed using a pipette and preserved in 5 ml of DNA preservation buffer (3.5 M ammonium sulphate, 17 mM sodium citrate, and 13 mM EDTA). Following collection, samples were concentrated by centrifugation at 3000× g for 15 ± 2 min at 5 ± 2 °C, frozen, and transported on dry ice to the UK Centre for Ecology & Hydrology (UKCEH), Wallingford, for subsequent analysis. Over a three-month period prior to biofilm sample collection, up to five water chemistry samples were taken, this data was used to calculate the water chemistry means.

Water quality measurements included nitrate, phosphorus, and dissolved oxygen levels, along with water temperature and pH were taken. Full water chemistry data are available in the supplementary material (Supp table 3, Supp. Fig 1).

### DNA extraction

DNA was extracted from 100 µL of biofilm suspension using the Quick-DNA Fecal/Soil Microbe Kit (Zymo Research, California, U.S.), with modifications to optimise the DNA yield (Newbold *et al.,* 2025). The extraction protocol was adapted as follows: 500 µL of DNA/RNA Shield (Zymo Research) was added to each sample as lysis buffer. The samples were mechanically disrupted using a TissueLyser II (Qiagen, Germany) at 20 Hz for 20 min. 20 µL of recombinant Proteinase K (Roche, Switzerland) was added to the lysate and incubated at 65°C for 20 min. The purified DNA was eluted in 100 µL of elution buffer. A negative extraction control, without sample material, was used to monitor potential contamination. DNA concentration was quantified using the Qubit dsDNA High Sensitivity kit (Life Technologies Limited) according to the manufacturer’s protocol. The extracted DNA was stored at 4 °C until PCR amplification. A detailed step-by-step extraction, PCR and library preparation protocol is available at https://doi.org/10.17504/protocols.io.j8nlk8em6l5r/v1.

### Sequencing

Two methods of sequencing were used to amplify the 16S rRNA gene (Fig. 1). Illumina NextSeq for short-read sequencing (291bp) and Pacific Biosciences Sequel II for long-read sequencing (1500bp).

**Figure 1:**
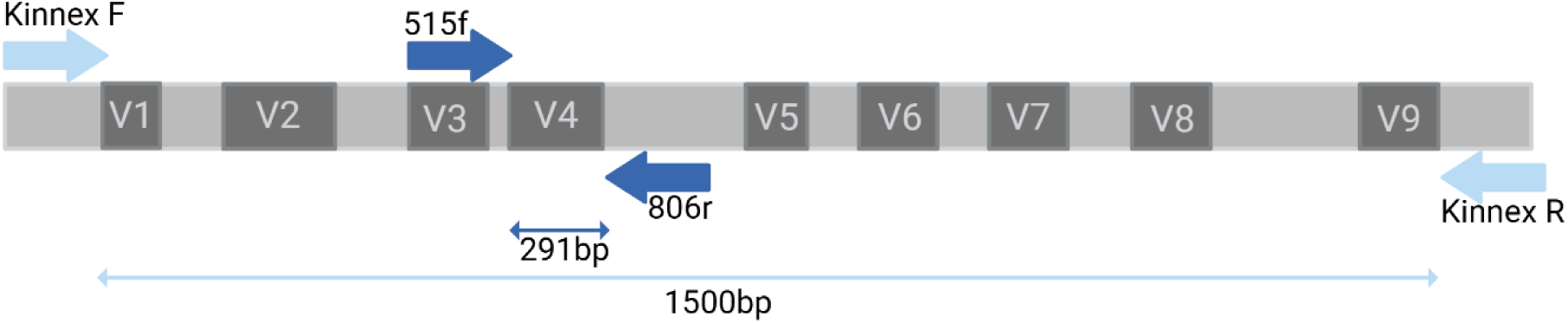
Primer positions on the 16S rRNA gene, showing overlap of the illumina 16SV4 primers within the 16S gene sequenced by the Kinnex primers.

#### Pacific Biosciences

The V1-V9 region of the 16S rRNA gene was amplified by Novogene using the Kinnex protocol (Srinivas *et al.,* 2025) (primer sequences in Table 1, PCR amplification in Table 2). The PCR products of the barcoded V1–V9 amplicons were detected by agarose gel electrophoresis prior to processing on a PacBio Sequel II sequencing platform (Novogene, Cambridge, UK).

#### Illumina

The V4 region of the 16S rRNA gene was amplified using specific primers (Table 1) modified to include Illumina adaptor sequences. In a UV-sterilized laminar flow hood, a master mix was prepared containing 0.5 µL of 2000 units mL^-1^ Q5 high-fidelity DNA polymerase, 10 µL of 5x reaction buffer, 10 µL of 5x high GC enhancer (New England Biolabs, UK), 1 µL of a 10 mM dNTP mix (Bioline, UK), 0.1 µL of each 100 µM forward and reverse primer pair (Table 1) and 26.3 µL of molecular grade water. The master mix (48 µL) was dispensed into each well of a 96-well plate, and 2 µL of template DNA was added per sample. Negative PCR controls were also included. The thermocycling conditions are presented in Table 2. Successful amplification was verified by 1.5% agarose gel electrophoresis using GelRed nucleic acid staining. PCR products were purified using a MultiScreen PCR filter plate, resulting in 35 µL of eluted product

The second PCR step employed a dual-indexing approach to enable sample demultiplexing. Indexing primers were prepared using an Opentrons liquid-handling robot, each consisting of a forward (i5) or reverse (i7) Illumina adaptor sequence, an i5 or i7 Nextera index and an Illumina pre-adaptor sequence. The second PCR mix contained 0.25 µL of Q5 DNA polymerase, 5 µL of reaction buffer, 5 µL of high GC enhancer, 0.5 µL of dNTPs, 5 µL of the indexing primers (pre-prepared in the plate), 7.25 µL of molecular grade water and 2 µL of purified PCR product from the first PCR step. The cycling protocol is presented in Table 2. Amplification was confirmed using agarose gel electrophoresis.

The second-step PCR product was normalised using the NGS Normalization kit (Norgen Biotek, Canada) to achieve a concentration of approximately 5 ng µL^-1^. The samples were pooled by plate and quantified using a Qubit High-Sensitivity Assay Kit. The amplicon library was prepared by diluting and pooling the samples, followed by concentration and purification using gel extraction. The final libraries were quantified, diluted to 1000 pM, and sent to Illumina Cambridge for sequencing on a NextSeq 2000 with a P1 flow cell and 40% PhiX control.

### Data Analysis

Amplicon sequence reads were processed using the DADA2 pipeline (Callahan *et al*., 2016) implemented in R [version 4.4.2]. Short-read and long-read sequences underwent distinct processing workflows, with full analysis scripts for short-read sequences available at https://github.com/amycthorpe/amplicon_seq_processing_biofilms. For short-read analyses, raw sequences were demultiplexed, and adaptor sequences were trimmed using the Illumina FASTQ generation pipeline. Primers were removed using the ’trimLeft’ parameter, and the quality profiles of the forward and reverse reads were examined. Reads were truncated when quality scores fell below Q30 and filtered using stringent criteria, including removing reads with ambiguous bases and a maximum expected error threshold of 2. The DADA2 algorithm learned error rates from a 100 million base subset, with visualisations confirming the alignment of the estimated rates with the observed data. Reads were then dereplicated into unique sequences based on the error rate model, and the core sample inference algorithm was used to identify true sequence variants. Paired forward and reverse reads were aligned and merged, requiring a minimum of 12-base overlap. Chimeric sequences were identified and removed, resulting in an amplicon sequence variant (ASV) abundance table. Long-read analyses followed a modified protocol (https://benjjneb.github.io/LRASManuscript/LRASms_fecal.html), with primers removed and reads trimmed to a minimum length of 1,200 bp and a quality threshold of three (filterAndTrim(nops2, filts2, minQ=3, minLen=1200, maxLen=1600, maxN=0, rm.phix=FALSE, maxEE=2)). Subsequent demultiplexing generated a sequence table with sample-specific counts. Taxonomy was assigned to each ASV using the naive Bayesian classifier (Wang *et al*., 2007) with a minimum bootstrap confidence of 60 against the SILVA v138.1 reference database (Quast *et al*., 2012) for Illumina and PacBio 16S rRNA gene sequences.

The sequences were rarefied to a uniform sequencing depth by examining rarefaction curves and identifying the sequencing depth at which the richness plateaued (Supp Fig. 6). Both long- and short-read data were rarefied to 3,000 reads per sample to conserve the majority of samples. All negative extraction and PCR controls and a small number of samples (five) did not meet the rarefaction depth and were therefore removed. Sequences assigned as mitochondria and chloroplasts were removed from the datasets. These accounted for 31% and 32% of the total reads in the short- and long-read data, respectively.

Downstream analyses were performed using R [version 4.4.2]. Counts, taxonomy, and metadata files were loaded into Microeco (Liu *et al*., 2021) for processing and visualisation. To assess differences in the number of ASVs between sequencing technologies, we used a Wilcoxon signed-rank test followed by a linear mixed effects model to evaluate the direction of the effect. Taxonomic assignment proportions across ranks were compared using the Mann– Whitney U test. Differences in the relative abundance of taxa between sequencing platforms were tested using a Wilcoxon rank-sum test with Bonferroni correction for multiple comparisons.

To examine community composition similarities between sample types and sequencing platforms, Principal Coordinates Analysis (PCoA) based on Bray–Curtis dissimilarity at the genus level was performed, along with a Procrustes analysis to assess concordance between ordinations. Prior to analysis, ASV identifiers were replaced with their corresponding taxonomic names from the kingdom to genus level to ensure meaningful comparison of community structure between sequencing methods. Additionally, Deming regression was used to investigate the agreement between the number of reads assigned to each phylum across sequencing types. All statistical analyses were performed in R, with significance thresholds set at p < 0.05.

To compare short- and long-read ASVs in the sequence space and identify overlapping short- and long-read ASVs, we used NCBI BLAST (Camacho *et al*., 2009), with the following parameters: ‘–id 80 –query-cover 90 –subject-cover 90 –more-sensitive –outfmt 100‘. The output was filtered to retain sequences with at least 90% identity and a minimum length of 235 bp for subsequent analysis. This length threshold was chosen as it represents 10% below the median short-read length. Short-read ASVs that did not map to long-read ASVs were considered as unique sequences.

To assess potential primer bias, the long-read sequences were trimmed using the specific primer sequences employed in the short-read protocol to isolate the corresponding amplicon region. These trimmed long-read sequences were then processed using the same DADA2 pipeline as the original long-read data. All subsequent analyses were conducted in R using the same workflow to ensure consistency and comparability.

The relevant code for the analysis and figures, and raw data can be found at: **10.5281/zenodo.15223954**.

## Results

### River Surveillance Network biofilms

At each sampling point (Fig. 2a), epilithic biofilms were collected to assess bacterial biodiversity, complemented by the concurrent collection of relevant water chemistry and nutrient data (Fig. 2b–e) to understand how biotic and abiotic factors shape the community composition of bacteria within the RSN. Water chemistry measurements, including pH, conductivity, and nutrient concentrations (e.g., nitrate, phosphate), showed variation across sites, with each parameter displaying a broad distribution (Supplementary Table 3). However, potential physicochemical drivers of sequencing technology differences were not further explored due to the limited number of sites (n = 7), which constrained our ability to draw statistical inferences.

**Figure 2:**
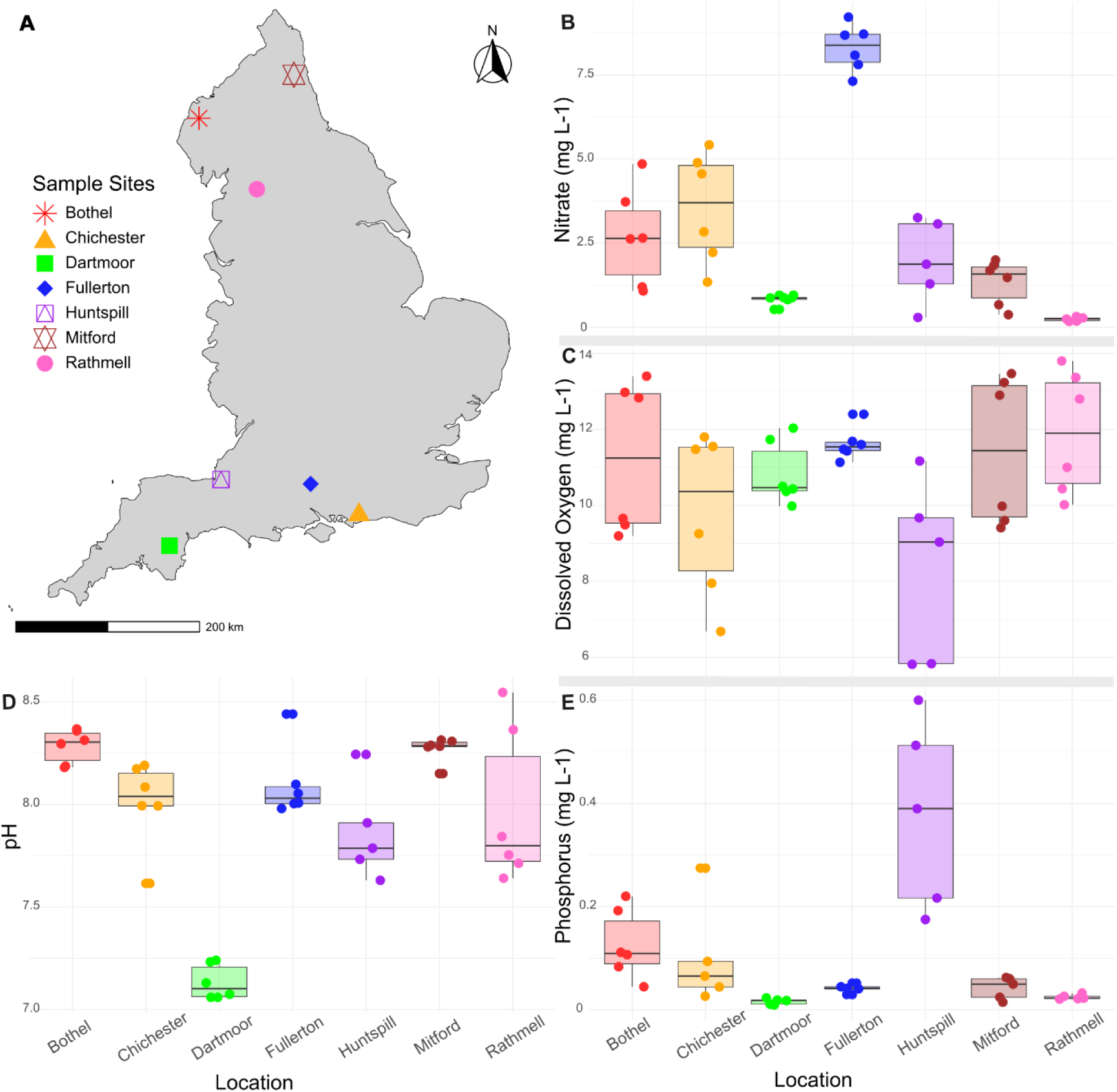
(A) Map of the seven sites across England from which the biofilm samples were collected. Water chemistry data from the seven sites; Nitrate (mg L^-1^) (B), dissolved oxygen (mg L^-1^) (C), pH (D), and Phosphorus (mg L^-1^) (E) levels at each sample point for each location. Water chemistry means were calculated using five chemistry samples collected over a three-month period prior to biofilm sample collection.

### Short- and long-read taxonomic compositions are similar

To compare the two sequencing approaches (short Illumina vs. long PacBio reads), we assessed community composition and the relative abundance of assigned taxa. While both methods recovered similar overall compositions at the phylum and genus levels, significant differences were observed in the relative abundance of several taxa based on a Wilcoxon rank-sum test with Bonferroni correction (Fig 3a-b). At the phylum level, Actinobacteriota (p=0.0005), Myxococcota (p<0.0001), Gemmatimonadota (p=0.00012), and Chloroflexi (p=0.014) were significantly more abundant in the short-read dataset. At the genus level, one notable difference was a significantly higher abundance of Ferruginibacter (p=0.0038) in the long-read dataset.

**Figure 3:**
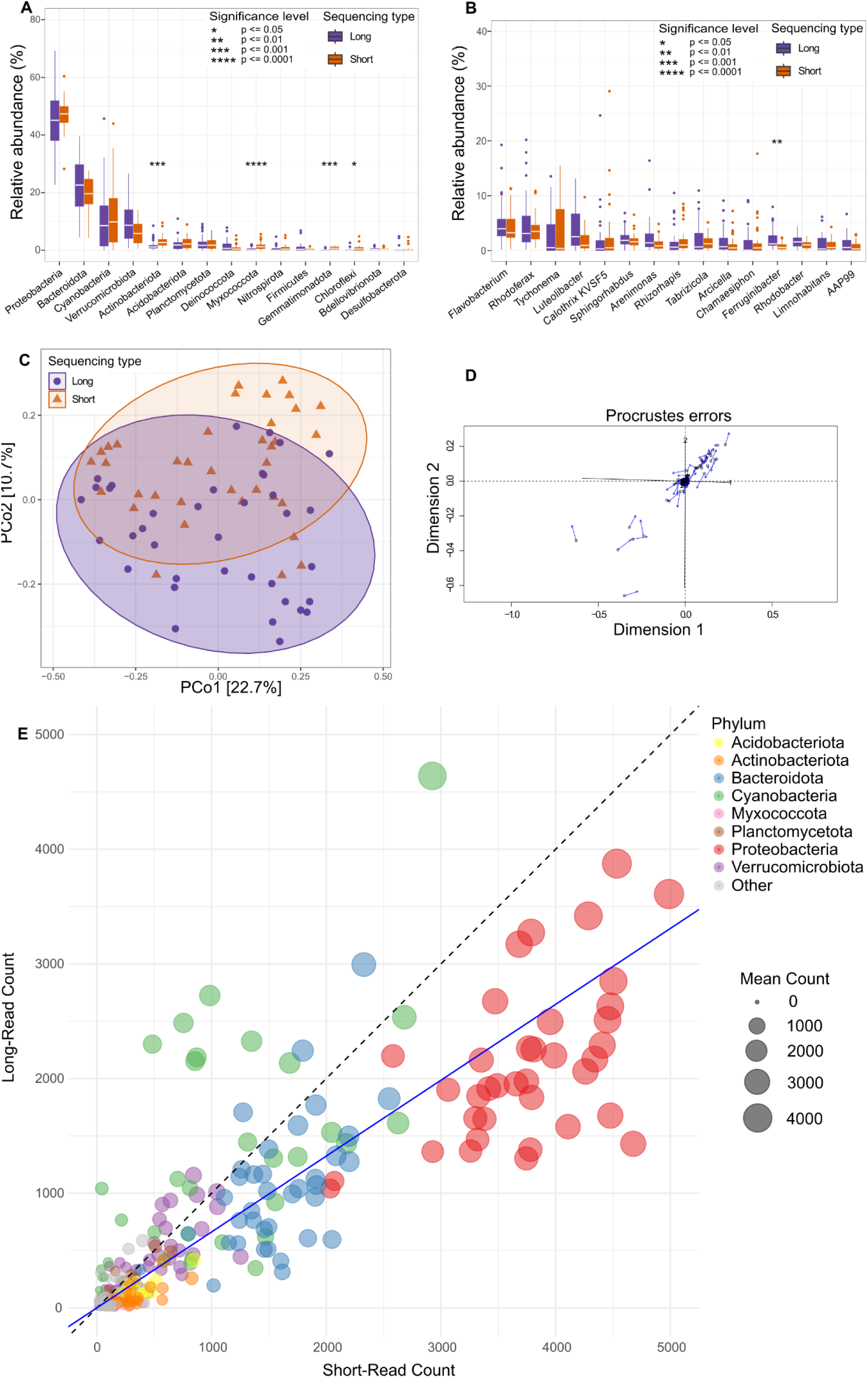
Comparison of bacterial taxon composition between short- (orange) and long-read (purple) sequences. Comparison of the top 15 abundant taxa at the phylum (A) and genus (B) levels for short- and long-read sequencing. Significant levels indicated by * show significant differences between long- and short-read relative abundances for each taxa. PCoA of paired samples for short- and long-read sequencing (C). (D) Procrustes error plot for paired short- and long-read samples (p < 0.001). (E) Scatter plot showing the relationship between short- and long-read abundance for each phylum across all the samples. The top eight phyla are colour-coded, and the circle size is proportional to the mean number of reads per phylum in each paired sample. The dashed black line represents the line of perfect fit (1:1), and the blue line depicts the Deming regression line, with a slope of 0.66.

An ordination analysis to assess the similarity between short-read and long-read results plot (Fig. 3c) showed partial overlap between the two sequencing methods, indicating shared community composition. To further quantify the similarity between the two datasets, a Procrustes analysis was conducted (Fig. 3d). The results showed a strong concordance between short-read and long-read sequencing compositions (R² = 0.91, p < 0.001), indicating that, despite methodological differences, both approaches captured comparable community structures. The Procrustes error plot visually represents the alignment between the datasets, with minimal deviation in most cases.

Furthermore, we assessed the number of reads assigned to bacterial phyla across sequencing methods to determine whether either approach exhibited taxonomic biases. We found a strong correlation (R2 = 0.66, Demming regression) between short-read and long-read sequencing abundances (Fig. 3e), but with a consistent bias towards higher read counts in the short-read dataset. The relationship between short and long-reads is based on a Deming regression analysis, which yielded a slope of 0.66 (95% confidence interval (CI): 0.60–0.74), indicating that, for most phyla, short-read sequencing recovered more reads than long-read sequencing.

However, phylum-specific trends were evident. Proteobacteria, for example, exhibited notably higher read counts in the short-read dataset, whereas Cyanobacteria were more relatively abundant in the long-read dataset. These deviations suggest that certain bacterial phyla may be differentially represented depending on the sequencing method, potentially due to differences in primer binding efficiency, amplified fragment length, error profiles, or taxonomic classification accuracy between methods.

### Improved taxonomic resolution with long reads

The number of ASVs was 68% higher in long-read sequencing (11,592 ASVs) than in short-read sequencing (7,985 ASVs) after rarefaction, the difference between pre- and post-rarefaction was not of note (Supp Fig. 5). At the per-sample level, paired analysis (Fig. 4a) revealed a significant difference in the number of ASVs produced by each sequencing method (Wilcoxon signed-rank test: V = 550.5, p = 0.0027), with the short-read method producing significantly more ASVs.

**Figure 4:**
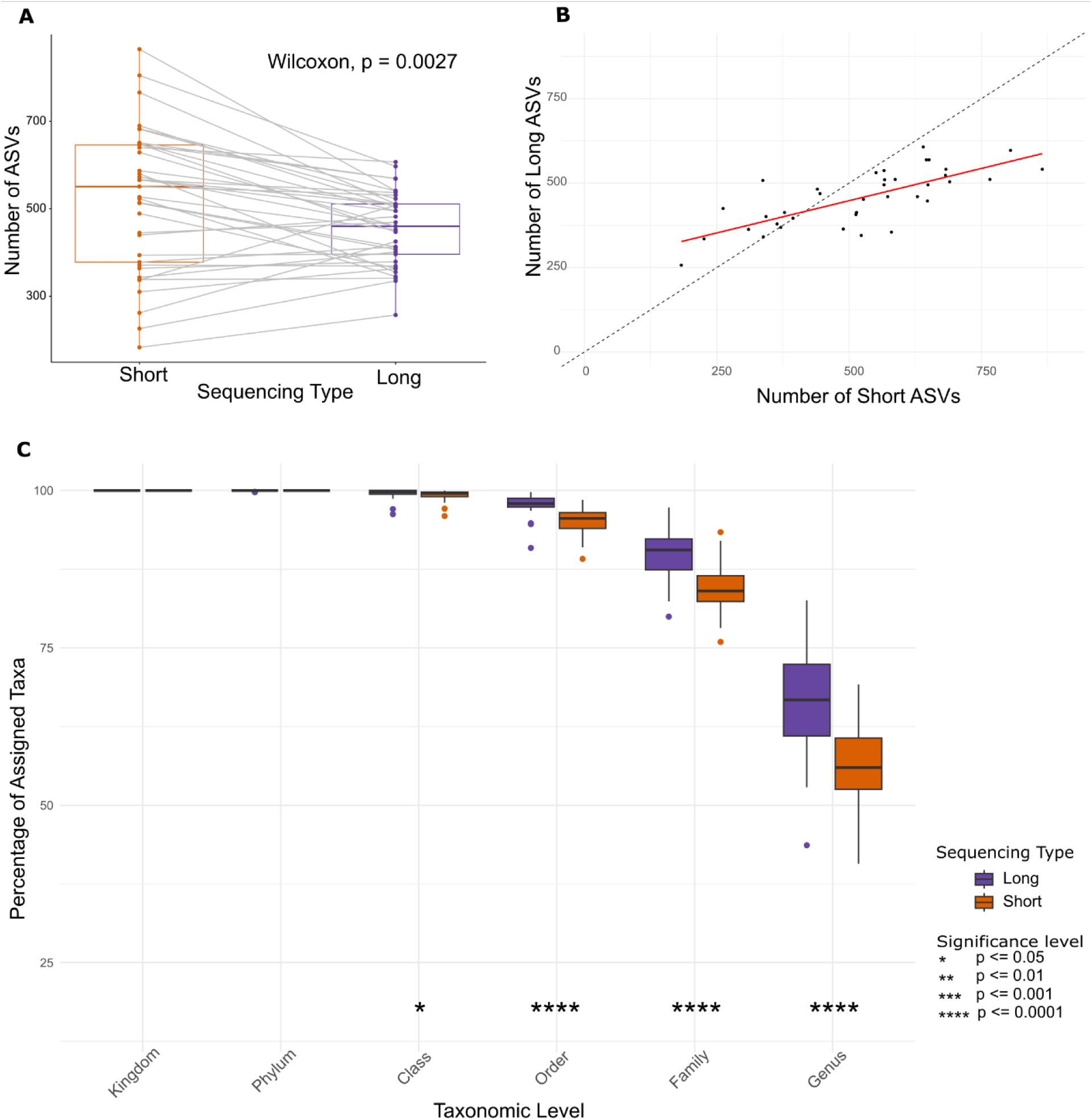
Comparison of taxonomic assignment and ASV detection between long-read (purple) and short-read (orange) sequencing methods. (A) Paired comparison of ASV counts per sample, analysed using the Wilcoxon signed-rank test (V = 550.5, p = 0.0027). (B) Relationship between ASV counts detected by short- and long-read sequencing, where the dashed black line represents a 1:1 ratio (perfect agreement), and the red line represents the fitted linear model (LLM). Short-read sequencing detected significantly more ASVs per sample than long-read sequencing, with an estimated increase of 65.38 ASVs (±18.94 SE, t = 3.45, p = 0.00144) per sample. (C) Percentage of assigned taxa at different taxonomic levels for long- and short-read sequencing, Mann-Whitney U test to test for significant difference between sequencing types.

**Figure 5:**
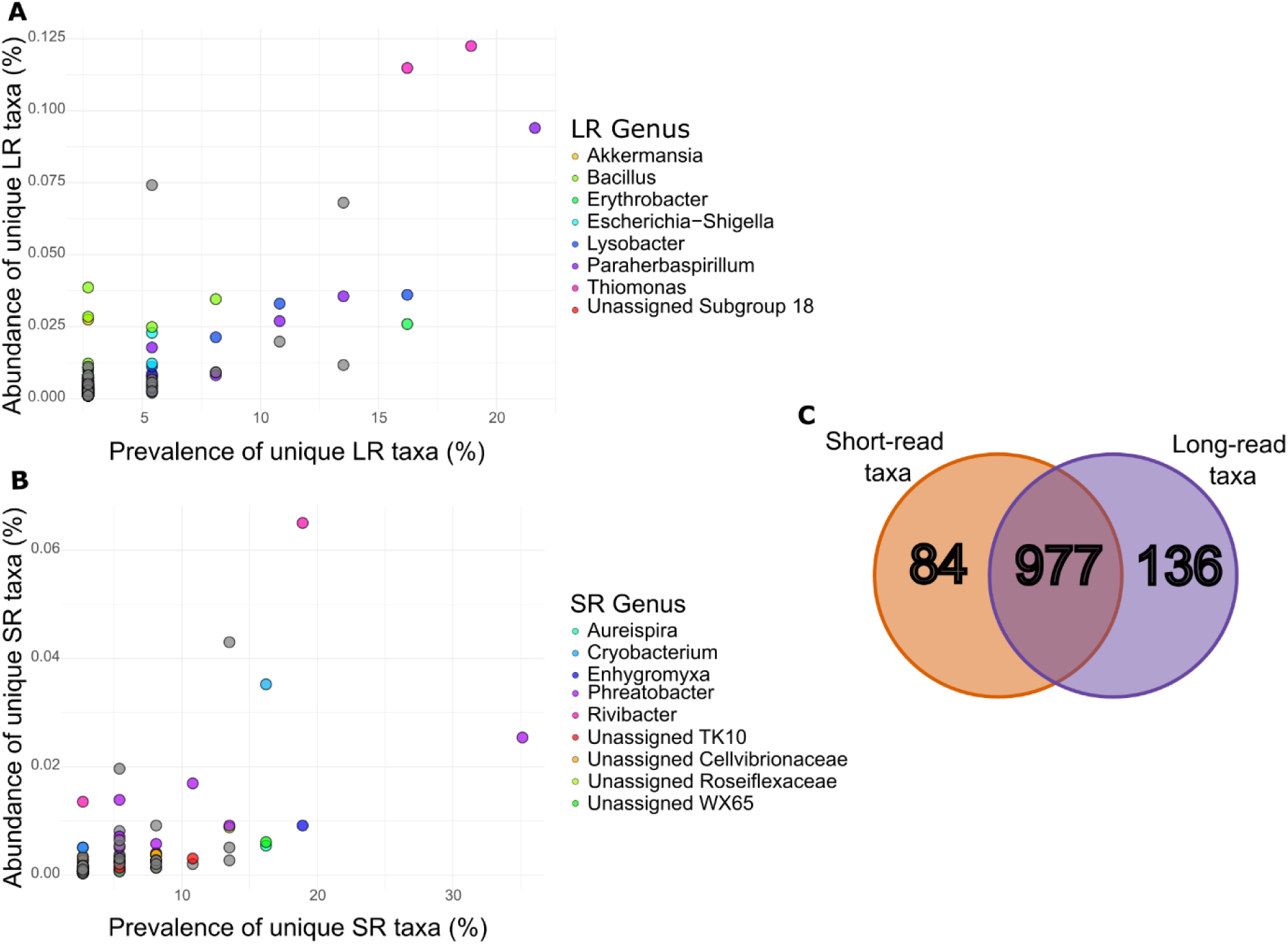
The prevalence (the % of total samples detected in) and abundance (the % of total reads detected in) of (A) long-read (LR) ASVs that did not have a >=98% match to any short-read ASVs in the whole dataset and (B) short-read (SR) ASVs that did not have a >=98% match to any long-read ASVs in the whole dataset, coloured by the top genera. (C) Venn diagram showing the number of unique taxa in the short-read (orange) and long-read (purple) data, and the overlap of the datasets.

A linear mixed-effects model (LMM) was used to assess the effect of sequencing type on the number of observed ASVs, while accounting for paired samples (Fig. 4b). The model included sequencing type as a fixed effect and sample identity as a random effect to account for variations among samples. The model showed that short-read sequencing detected significantly more ASVs than long-read sequencing, with an estimated increase of 65.38 ASVs (±18.94 SE, t = 3.45, p = 0.00144) per sample. The random effect of sample identity had a variance of 10,155 (SD = 100.77), indicating a substantial between-sample variability. The residual variance was 6,636 (SD = 81.46), reflecting the within-sample variation after accounting for sequencing type.

We assessed the proportion of taxonomic assignments across various ranks to determine whether the sequencing type affected taxonomic resolution. At both the kingdom and phylum levels, there were no significant differences in the percentage of taxa assigned between the short- and long-read sequencing methods; 100% of the taxa were assigned at these ranks using both approaches. At the class level, 98.9% of the taxa were assigned using short-read sequencing, compared to 99.6% with long-read sequencing—a small but statistically significant difference (p = 0.02; Mann–Whitney U test). More pronounced differences were observed at lower taxonomic ranks. Long-read sequencing resulted in significantly higher proportions of taxonomic assignment at the order (4.4% increase, p < 0.0001), family (5.8% increase, p < 0.0001), and genus (11.6% increase, p < 0.0001) levels compared to the short-read method (Fig. 4c).

### Unique features of sequencing type

Given the substantial taxonomic overlap between the sequencing methods, we evaluated the sequence similarity of the short- and long-read ASVs to determine which were truly unique. Using NCBI BLAST (Camacho et al., 2009), we found that on average 83.8% (58%-96%) of short-read ASVs aligned with long-read ASVs (Supp. Fig. 4a) within the expected 350–750 bp region of the long-read 16S sequences, corresponding to the V4 primers used for short-read amplicon sequencing (Supp. Fig. 3).

To test whether sequence identity cutoffs in BLAST influence mapping of short-read ASVs to long-read sequences, we examined the alignments at different identity thresholds. At a relaxed identity cutoff (90%), over 95% of short-read ASVs had a corresponding long-read match, whereas at a stringent 100% identity cutoff, this proportion dropped to ∼60% (Supp. Fig. 4a). Notably, short-read ASV length (230 bp vs. 253 bp) did not affect mapping success, as both lengths exhibited comparable match percentages (Supp. Fig. 4c). These lengths were selected based on the minimum and median short-read ASV lengths.

Importantly, we analysed the composition of unique short-read ASVs that had no matches in the long-read dataset and found that they were distributed across multiple phyla. Interestingly, these short-read ASVs were predominantly associated with taxa of low prevalence (percentage found in all samples, average 5.6%) and abundance (percentage occurs in total ASVs, average 0.004%) (Fig. 5b). The reverse was also true for unique long-read ASVs that had no matches to the short-read dataset. These long-read ASVs are mainly associated with taxa of low prevalence (average 3.5%) and abundance (average 0.006%) (Fig. 5a).

A comparison of taxonomic annotations revealed that 977 taxa were shared between the short-read and long-read datasets, while 84 taxa (∼7% of the whole dataset) were unique to short-read sequencing and 136 taxa (∼11% of the whole dataset) were unique to long-read sequencing (Fig. 5c). These counts were based on full taxonomic annotations from each dataset.

## Discussion

The dynamic spatial and temporal complexity of river ecosystems creates habitats that drive the remarkable diversity and ecological richness of aquatic microbial communities. Epilithic river biofilms represent intricate ecological niches where many environmental parameters may influence the structure and composition of microbial communities (Shibabaw *et al*., 2021). Consequently, the accurate characterisation of these microbial communities requires molecular approaches that can capture the subtle taxonomic and functional diversity inherent in these dynamic systems. To comprehensively investigate the microbial landscape across seven distinct river sites in England, we used two sequencing technologies, Illumina short-read and PacBio long-read sequencing, targeting the 16S ribosomal RNA (rRNA) gene. This approach enabled a comparative assessment of the microbial community structure, providing the opportunity to distinguish the capabilities of each sequencing platform in resolving complex bacterial assemblages embedded within river biofilm environments.

Our analysis revealed that, for 16S rRNA-based taxonomic assessments of river biofilms, the choice of sequencing method (Illumina short-read or PacBio long-read) did not significantly influence the relative abundance of taxa within bacterial communities. Despite the disparity in read length and taxonomic resolution, both sequencing platforms produced broadly comparable abundance profiles across major taxonomic groups. This suggests that short-read sequencing, although limited in its ability to resolve taxa at deeper levels, still captures reliable patterns in community structure. Consequently, it remains a practical and informative approach for studies focused on broad-scale surveys or relative abundance patterns. These findings are consistent with previous studies (e.g., Butt et al., 2022; Buetas et al., 2024), which reported largely concordant diversity measures between sequencing platforms, and support the use of either method depending on the specific ecological question being asked.

Although long-read sequencing improved taxonomic resolution, particularly at finer levels, it did not significantly alter per-sample richness or diversity estimates (e.g., Chao1 or Shannon; Supp. Fig. 2). It must however be noted that the environmental gradients are the likely selecting factors reflected in the procrutes analysis which may be a stronger driver compared to technical variabilities in read length. Interestingly, while the total number of unique ASVs across the dataset was higher in the long-read data, individual samples contained significantly more ASVs in the short-read data. This likely reflects greater sequencing depth in the short-read dataset, allowing detection of more low-abundance variants per sample. In contrast, the higher resolution of long-read data can split similar sequences into more distinct ASVs across the dataset, inflating the total count. This inflation of total ASV count in the long-read data highlights how sequencing platform choices must be considered when interpreting ASV-based metrics.

Importantly, our findings demonstrate that long-read sequencing provides superior taxonomic resolution compared to short-read methods for analysing microbial communities within river biofilms. This aligns with a report by Gao *et al*. (2024), who also reported an increased taxonomic resolution of ASVs, including precision at the species and strain levels. However, we did not perform species-level taxonomic comparisons, as only a small proportion of ASVs could be confidently assigned to species across both sequencing methods. This limited resolution likely stems from incomplete reference databases for freshwater microbes, combined with the challenges of accurate species-level classification using current algorithms, especially for short-read data. Consequently, we focused our comparisons on higher taxonomic ranks where assignments were more robust and informative.

PacBio’s ability to generate longer contiguous sequences facilitated a more accurate taxon classification, particularly at deeper taxonomic levels, such as genera and species. This was also recently shown by Buetas *et al*. (2024), who found that all species were correctly identified using PacBio sequencing compared to Illumina short reads using a mock community. This advantage is particularly evident in complex microbial assemblages, where short-read sequences often fail to resolve ambiguities owing to their limited length and reliance on overlapping fragments for their assembly. To retrieve as much taxonomic information as possible we used lower bootstrap values which may have contributed to inflated assignment rates. Irrespectively, our analyses revealed that many sequences that remained unclassified in the short-read data matched successfully identified taxa in the long-read dataset. This improved classification makes the long-read dataset inherently more robust and reliable for the comprehensive assessment of biodiversity. This increase in the number of classified taxa is important for biodiversity assessments, as it allows for a more comprehensive understanding of community composition and structure. Such detailed insights are essential for applications such as biomonitoring and ecological research, where understanding the full spectrum of biodiversity is a priority.

Both methodologies, as highlighted previously by existing research, have advantages and disadvantages (Buetas *et al*., 2024; Eisenhofer *et al*., 2024; Gao *et al*., 2024). For example, Buetas *et al*. (2024) highlighted that Illumina provided an 8-fold higher throughput and lower cost than PacBio. Although in their study and ours, we identified a higher number of ASVs at the per sample level in the Illumina data, the overall increase in ASVs was not exponential in relation to the cost. This was similarly reported by Cook *et al*., who reiterated the need for deeper sequencing with long-reads to achieve parity with the short-read methodology. This is further supported by the observation of unique taxa within the Illumina samples compared to the PacBio samples in our study. Similar to the findings of Buetas et al. (2024), the unique ASVs identified by the short-read method were of relatively low abundance (∼0.004%) and prevalence (∼5.6%). We found the same trend in the long-read dataset, where unique ASVs had an average abundance of ∼3.5% and prevalence of just ∼0.006%. In terms of taxonomic assignments, 84 taxa were unique to short-read sequencing, 136 were unique to long-read sequencing, and 977 were shared between both methods. These unique taxa represented a small proportion of the total detected diversity—approximately 7% for short-read and 11% for long-read data. The higher number of unique taxa in the long-read dataset likely reflects the increased sequence length, which enhances the ability to resolve subtle differences between closely related organisms. Despite these differences, the substantial overlap in taxonomic composition demonstrates that both methods capture broadly consistent community profiles. From a biomonitoring perspective, this highlights the utility of long-read sequencing for enhancing taxonomic resolution without compromising comparability with established short-read approaches. Based on our findings, even at the genus level, there was a large inconsistency between the PacBio and Illumina taxonomic assignments (Supp. Fig 4b). This is likely due to the current state of databases, which are varied and not yet fully standardised or validated. For example, bespoke databases were developed by Lo *et al*. (2023) for aquatic pathogens. Similarly, others have reported that despite PacBio sequencing annotating more reads to the species level, the vast majority were taxonomically unassigned because of the possible under-representation of species in databases (Pasolli *et al*., 2019). With the increase in long-read methods for biomonitoring, database selection will have a critical impact. Our findings are in concordance with other reports, as highlighted above; therefore, the availability of curated databases in the future will play a major role (Sierra *et al*., 2020).

Collectively, our findings comparing the utility of short- and long-read methods for biomonitoring across the RSN revealed the accuracy and possible pitfalls of each sequencing technology. We acknowledge several limitations exist, including but not limited to primer bias, differences in the amplified 16S rRNA gene regions, and potential error/correction rates across the two methodologies. Primer bias, in particular, can affect which taxa are preferentially amplified, potentially skewing community composition. To mitigate this, we trimmed the long-read sequences to match the same region amplified by the short-read primers and re-analysed the data using the DADA2 pipeline with the same downstream processing, where ordinations revealed that the trimmed long reads overlapped across both the original long-read dataset and the short-read dataset. Furthermore, taxonomic assignment and ASV richness in the trimmed dataset were similar to those from the short-read data (Supplementary Fig. 7). These findings confirm that primer bias and the sequenced regions potentially contribute to observed differences in community composition. However, this bias is an expected feature of amplicon sequencing, and our study was specifically designed to assess the impact of sequencing technology, rather than primer performance, on taxonomic resolution and community profiling. Similarly the existing workflows pertaining to error corrections are still in their infancy with long reads and these areas require future research to understand the nuances of user choices in influencing outcomes of eDNA analysis (Bylemans *et al.,* 2025).

Despite this, we showed an increased resolution of taxonomic assignment using PacBio long-read sequence data, while simultaneously highlighting the possible inadequacy of sequencing depth using this platform, which can currently be achieved more cost-effectively using short-read technologies such as Illumina. It is important to reiterate that the optimal sequencing strategy depends on the research question, and that the specific goals or priorities of the study may dictate which method is chosen as appropriate. Overall, our data provide critical insights into the current molecular biomonitoring landscape and may serve as a valuable resource for future comparisons and subsequent benchmarks, particularly in environmental and ecological contexts.

## Acknowledgements

Funding

M.A. is supported by NERC SCENARIO PhD studentship [NE/S007261/1]. This work was supported by the Environment Agency under research project SC220034. S.B.B. is supported by the BBSRC Institute Strategic Programme: Decoding Biodiversity (DECODE) grant. A.C.T., J.W., K.W. and D.S.R. are supported by the Natural Environment Research Council (NERC) [grant numbers NE/X015947/1 and NE/X015777/1].

The views expressed in this paper are the views of the authors and do not represent the position of the Environment Agency or other employer organisations.

## Authors contribution

Physicochemical data and samples were collected by the Environment Agency. D.S.R., M.A., K.W., and J.W. conceptualised the study. M.A. and A.C.T performed sample processing, DNA extractions and sequencing. M.A. and S.B.B. designed and performed the bioinformatic analysis. M.A. performed statistical analysis and figure production. M.A. and S.B.B. drafted the manuscript. All authors read and revised the manuscript.

## Data Accessibility statement

Raw sequence reads and related metadata can be found here: **10.5281/zenodo.15223954**

