## Supplementary material for "Unlocking River Biofilm Microbial Diversity: A Comparative Analysis of Sequencing Technologies"

Supplementary Table 1: Primers used in this study.

| Sequencing Method | Forward primer (5’-3’) | Reverse primer (5’-3’) | Reference |
| --- | --- | --- | --- |
| Illumina 1st round | 16S rRNA 515f:  GTGYCAGCMGCCGCGGTAA | 16S rRNA 806r: GGACTACNVGGGTWTCTAAT | Walters et al. 2016 |
| Illumina 1st round adaptor | Forward adaptor: TCGTCGGCAGCGTCAGATGTGTATAAGAGAC | Reverse adaptor: GTCTCGTGGGCTCGGAGATGTGTATAAGAGACAG |  |
| Pacific Biosciences | Kinnex 16S F:  AGRGTTYGATYMTGGCTCAG | Kinnex 16S R:  RGYTACCTTGTTACGACTT | Kinnex V1-V9 |

Supplementary Table 2: PCR conditions.

| Sequencing Method | Stage | Temperature °C | Time | Number of cycles |
| --- | --- | --- | --- | --- |
| Illumina 1st round | Initial denaturation  Denaturation  Annealing  Extension  Final extension | 95  95  50  72  72 | 2 min  15 sec  30 sec  30 sec  10 min | 30 |
| Illumina  Indexing | Initial denaturation  Denaturation  Annealing  Extension  Final extension | 95  95  50  72  72 | 2 min  15 sec  30 sec  30 sec  10 min | 8 |
| Pacific Biosciences | Initial denaturation  Denaturation  Annealing  Extension  Final extension | 95  98  57  72  72 | 3 min  20 sec  30 sec  75 sec  5 min | 20 |

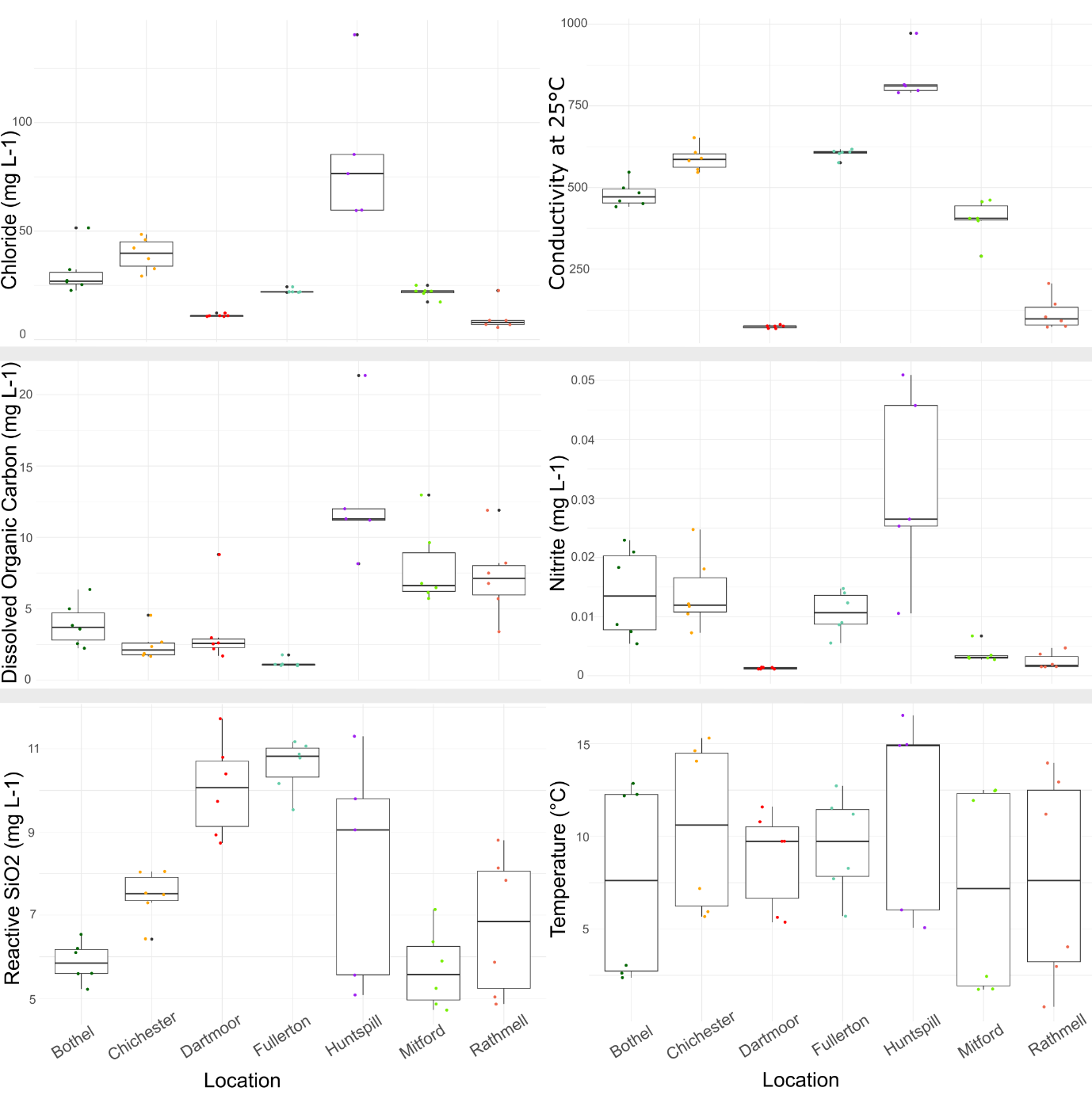

Supp Figure 1: chemistry data for the seven sampling sites; Chloride, Conductivity, Dissolved Organic Carbon, Nitrite, Reactive SiO2, and Temperature.

Supp Figure 2: Observed, Chao1 and Shannon Alpha diversity index for short-read and long-read sequencing. Shown as box plots and violin plots.

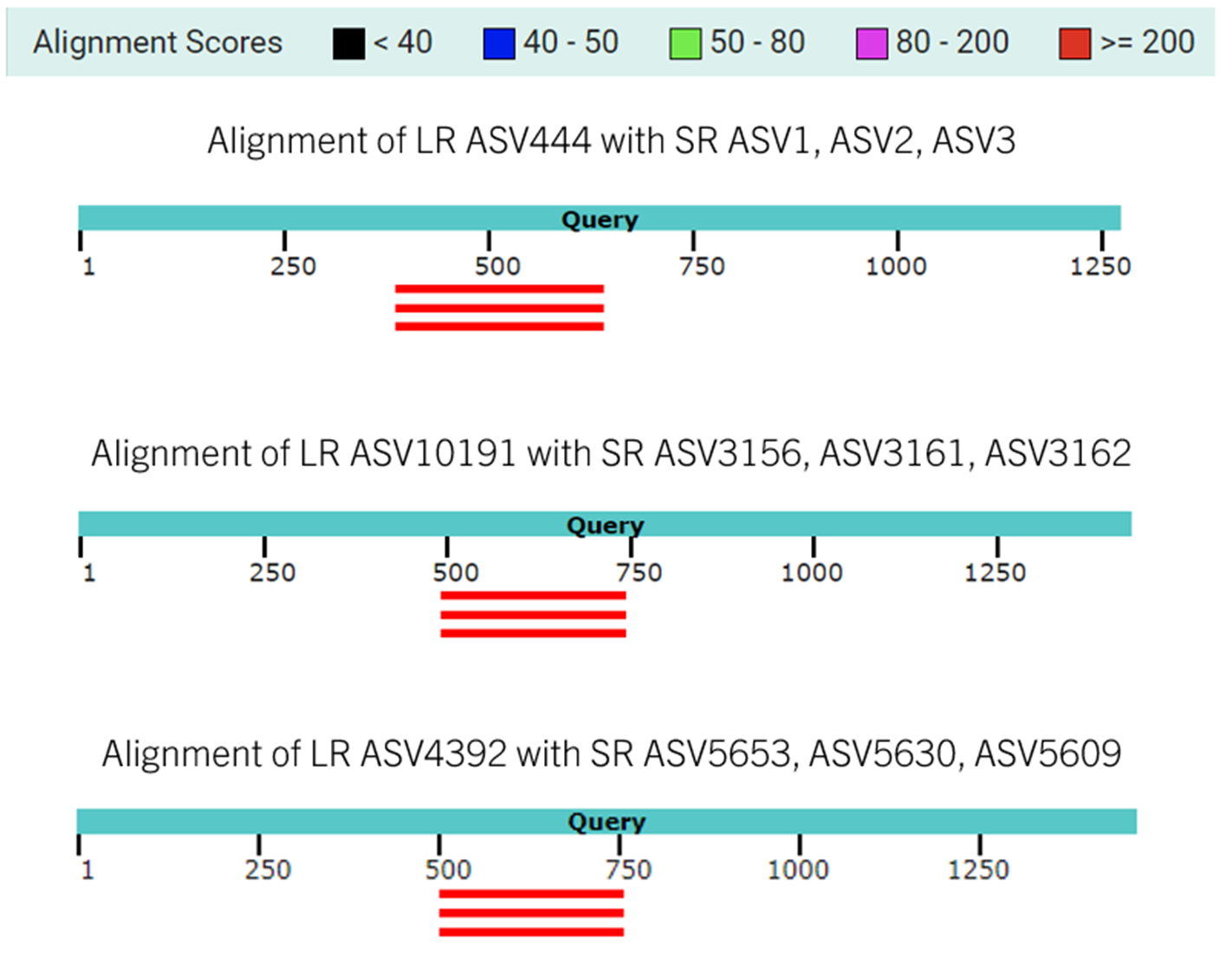

Supp Figure 3: Visualisation of BLAST alignments between long-read (LR) and short-read (SR) ASVs. The blue bar represents the LR ASV query sequence, while the red bars indicate the aligned SR ASVs that match the LR sequence. Alignment scores are colour-coded along the top bar, with red indicating a stronger alignment.

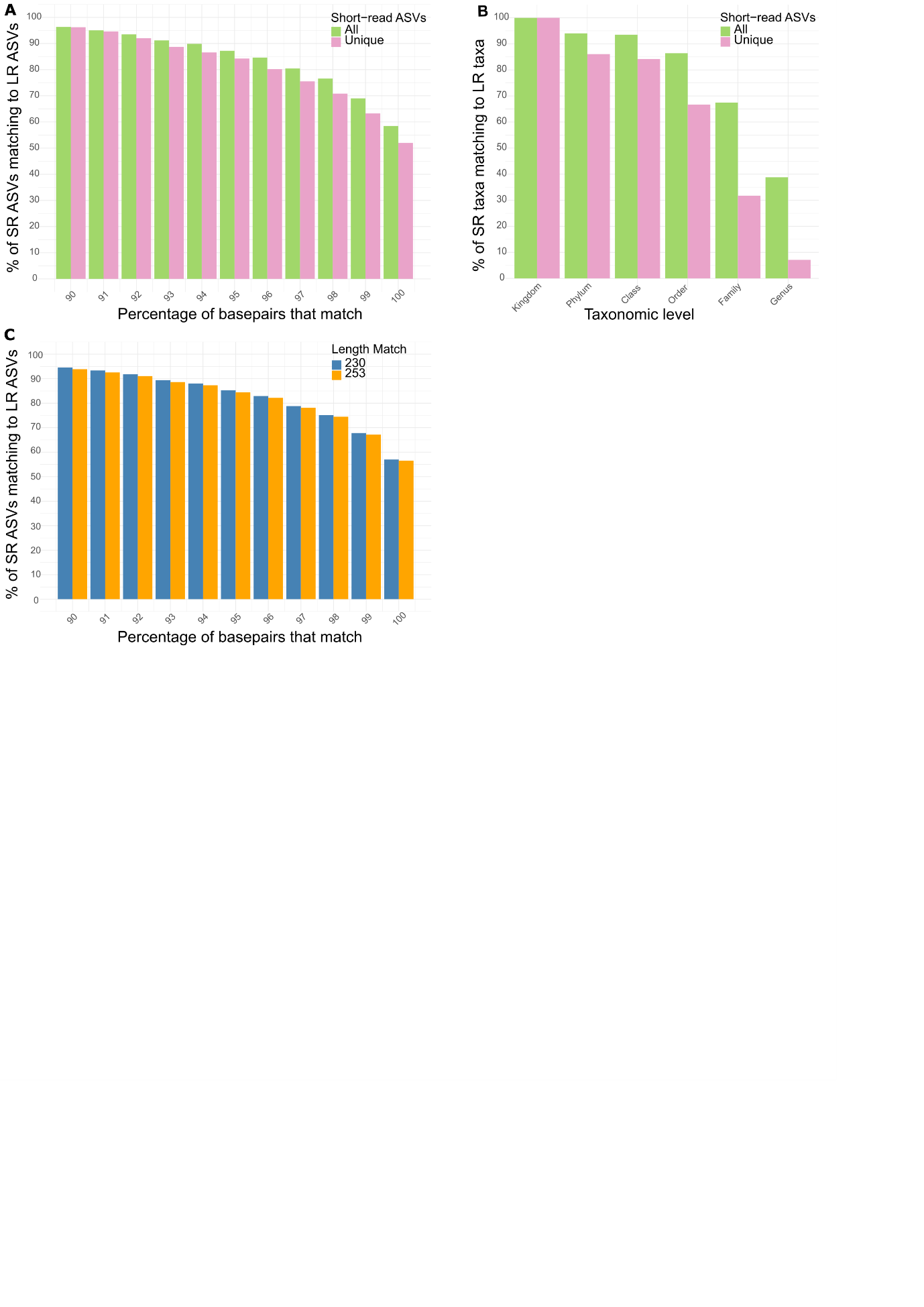

Supp Figure 4: Comparison of short-read (SR) ASVs mapped to long-read (LR) ASVs. (A) Percentage of short-read ASVs matching long-read ASVs across different sequence identity thresholds, with "All” SR ASVs in green and "Unique" SR ASVs in pink. (B) Taxonomic overlap of short-read ASVs mapped to long-read ASVs at different classification levels. (C) Effect of short-read ASV length (230 bp vs. 253 bp) on the matching probability to long-read ASVs.

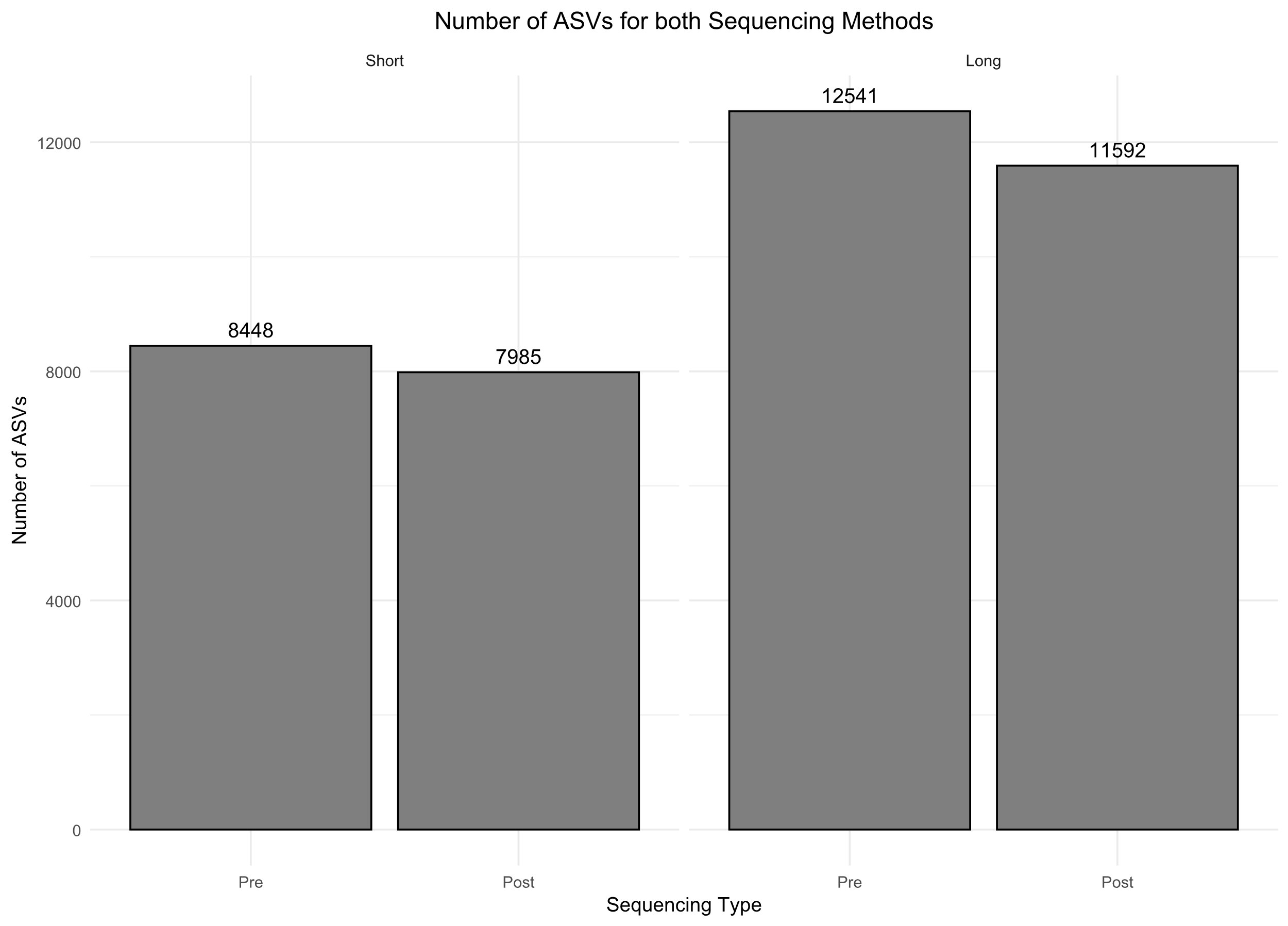

Supp Figure 5: The number of ASVs pre and post rarefaction for short-read and long-read sequencing.

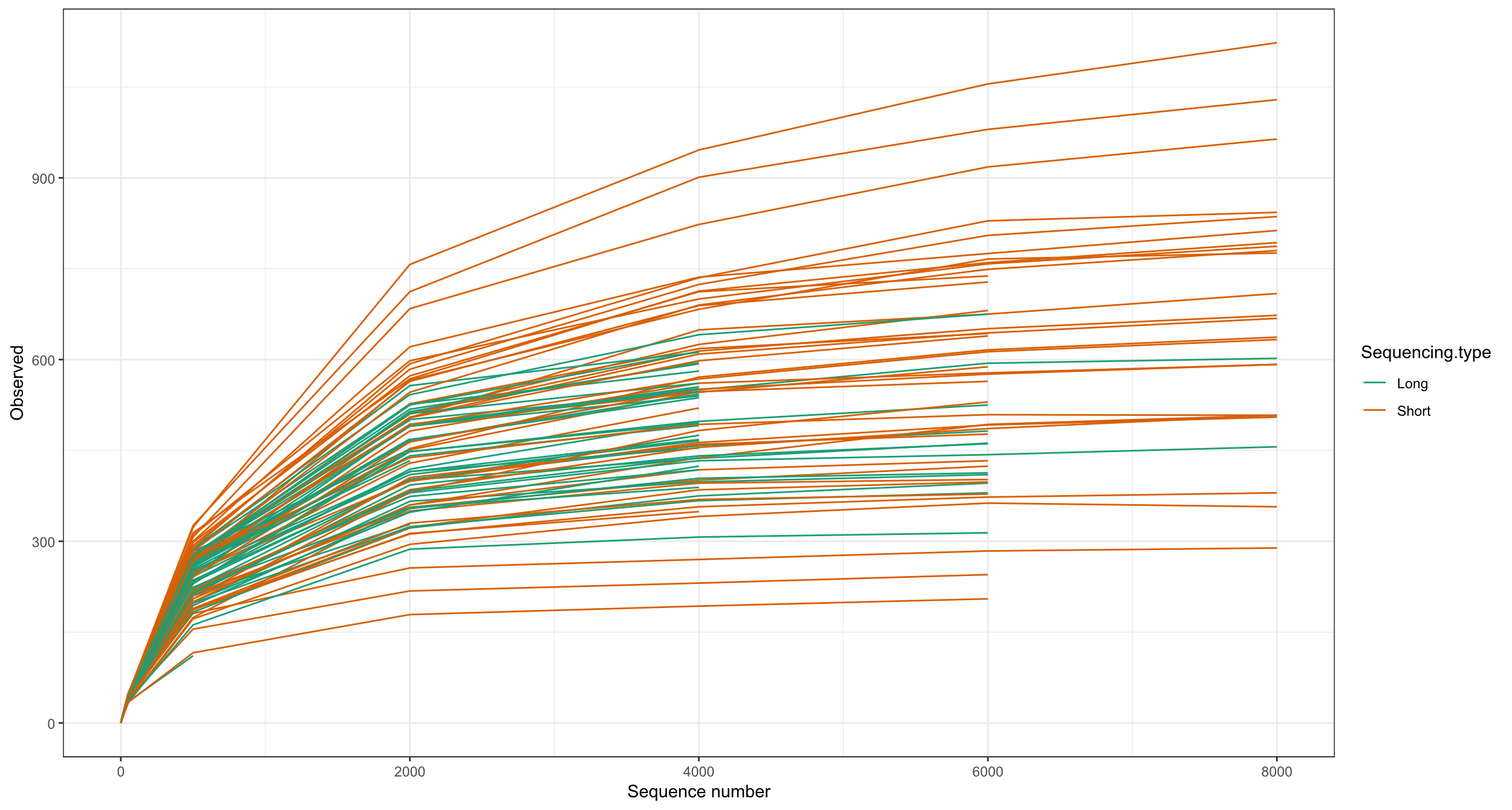

Supp Figure 6: Rarefaction curve for both long-read and short-read samples.

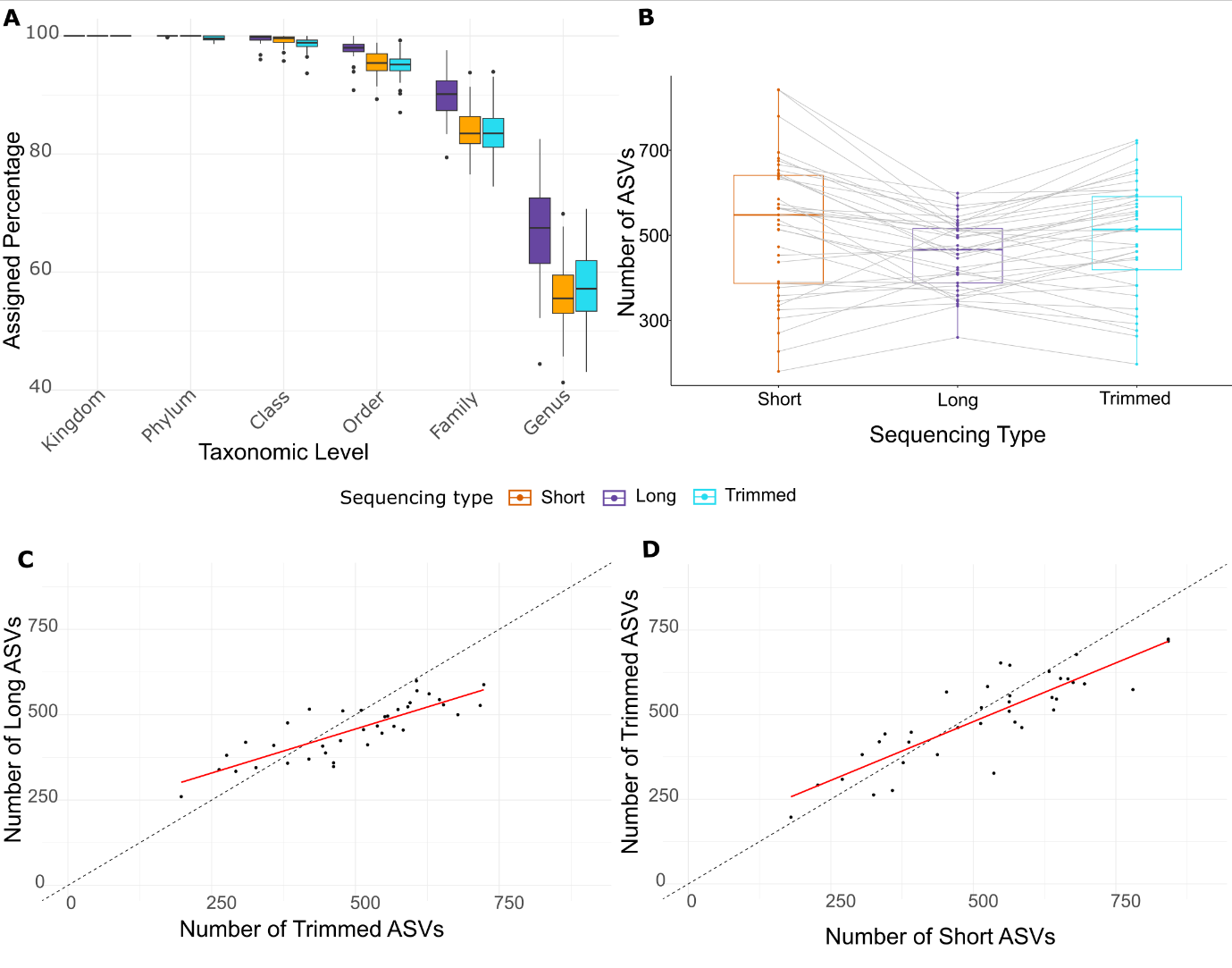

Supp Figure 7: Comparison of taxonomic assignment and ASV detection between long-read (purple), short-read (orange) and trimmed long-read (blue) sequences. (A) Percentage of assigned taxa at different taxonomic levels for long-, trimmed long- and short-read sequencing. (B) Paired comparison of ASV counts per sample. Relationship between ASV counts detected by long-read and trimmed long-read (C), and short-read and trimmed long-read (D) sequences, where the dashed black line represents a 1:1 ratio (perfect agreement), and the red line represents the fitted linear model (LLM).

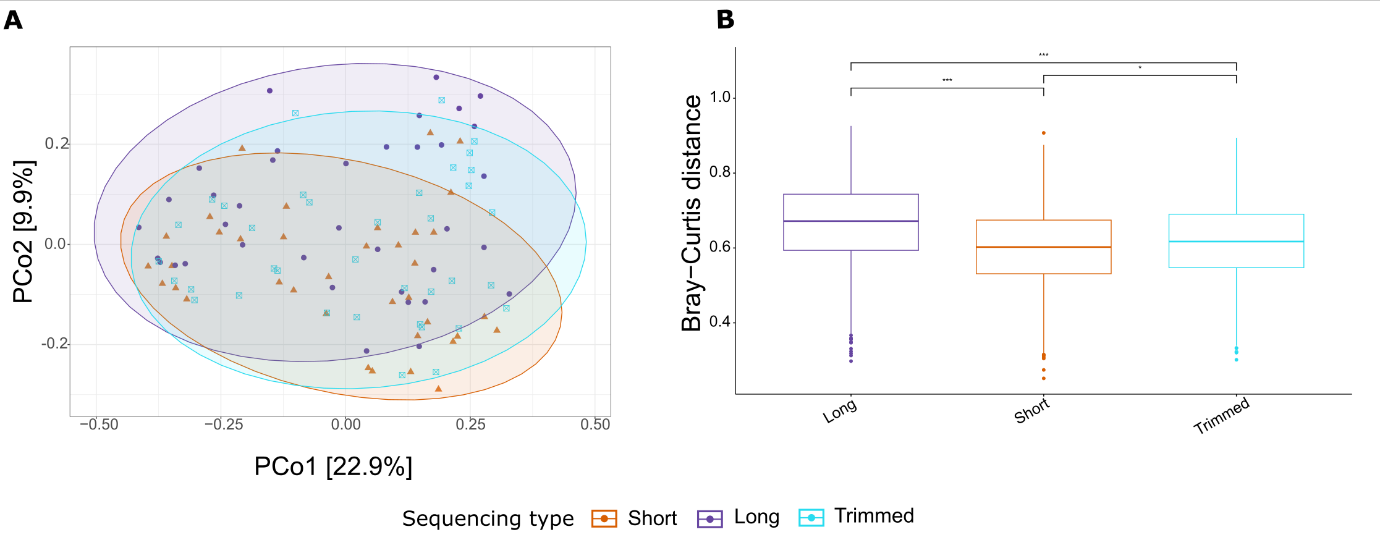

Supp Figure 8: PCoA of paired samples for short- (orange), long-read (purple) and trimmed long-read (blue) sequences (A). Bray-Curtis distance plot for the same samples (B)

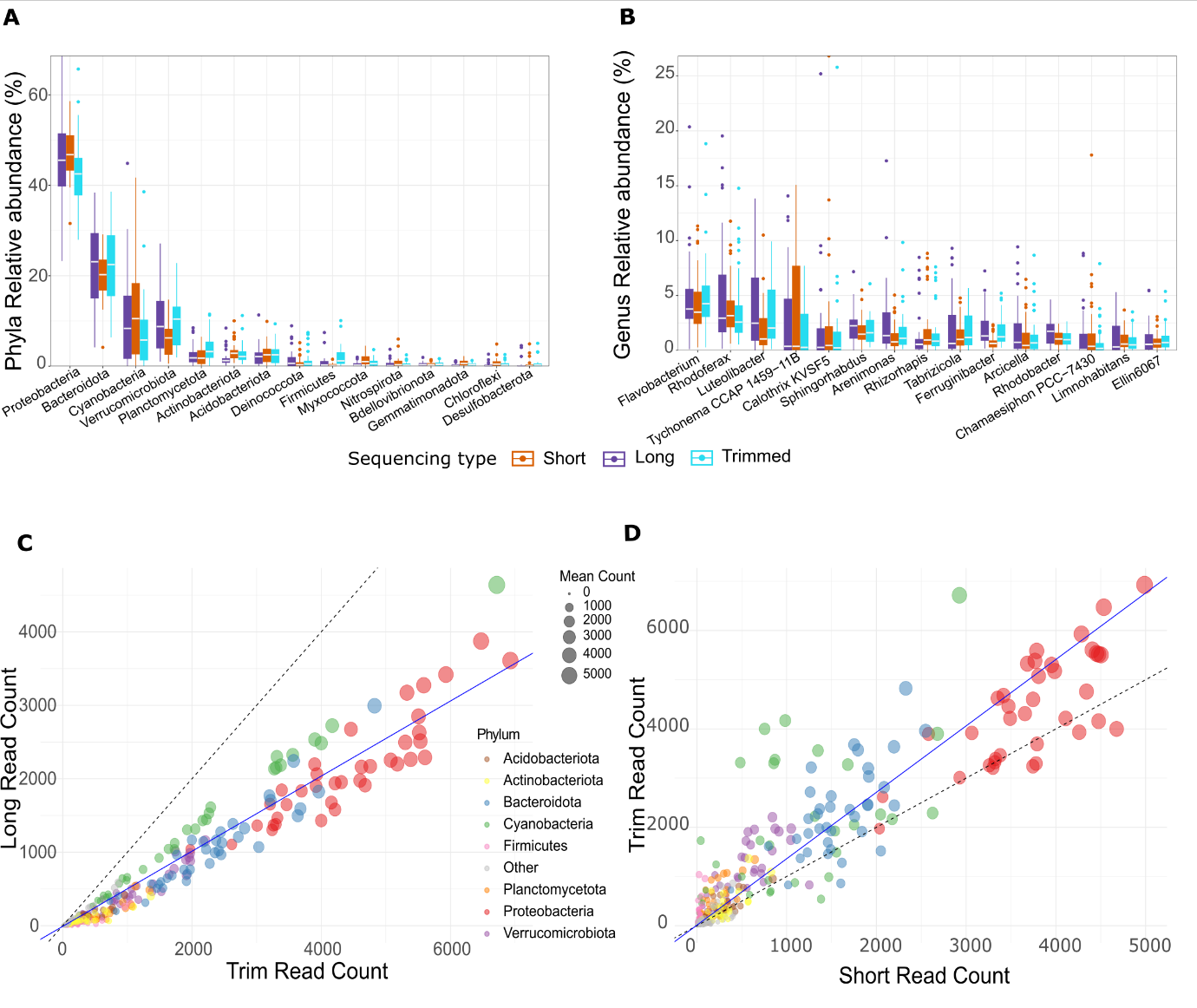

Supp Figure 9: Comparison of bacterial taxon composition between short- (orange), long-read (purple) and trimmed long-read (blue) sequences. Comparison of the top 15 abundant taxa at the phylum (A) and genus (B) levels for short-, long-, and trimmed-reads. Scatter plot showing the relationship between long-read and trimmed long read (C) and short-read and trimmed long-read (D)abundance for each phylum across all the samples. The top eight phyla are colour-coded, and the circle size is proportional to the mean number of reads per phylum in each paired sample. The dashed black line represents the line of perfect fit (1:1), and the blue line depicts the Deming regression line, with a slope of 0.51 (C) and 1.35 (D).
